# Geocoding genomic databases using GBIF

**DOI:** 10.1101/469650

**Authors:** Roderic D. M. Page

## Abstract

Many nucleotide sequences in the publicly available genomics databases lack spatial information, such as the latitude and longitude coordinates for the locality where the sample for sequencing was taken. In this note I discuss several approaches to geocoding sequence records. The first method uses the Global Biodiversity Information Facility (GBIF: https://gbif.org) as a gazetter. The availability of a simple full text search across GBIF data makes it possible to rapidly geocode locality information simply by searching for matching records within GBIF. Hence if a sequence lacks coordinates but has some locality information it could be rapidly geocoded. The second method matches voucher specimen code for sequences with the corresponding specimen records in GBIF, which may be geocoded even if the sequence obtained from that specimen is not. Lastly, there will be cases where sequence records lack either locality or specimen information, but that information is available elsewhere, such as in the published literature or in supplementary data files. The possibility of publishing geocoded sequence records using Github is discussed.

## Motivation

The DNA Data Bank of Japan (DDBJ), the European Nucleotide Archive (ENA), and GenBank together comprise the International Nucleotide Sequence Database Collaboration (INSDC) (Karsch-Mizrachi, et al., 2018). Numerous sequences in the INSDC databases have latitude and longitude information, but the majority do not (Marques et al. 2013, Schindel et al. 2013). Several researchers have developed methods to geocode sequences (Gratton et al., 2016; Miraldo et al., 2016; Tahsin et al., 2017; Weissenbacher et al., 2015), which often makes use of data sources such as GeoNames (https://geonames.org). However this approach overlooks probably the single largest database of geocoded biological specimens, namely the Global Biodiversity Information Facility (GBIF: https://gbif.org). This note outlines ways GBIF could be used to help geocode genomic information, and extends this to include additional, literature-based methods.

For large-scale geocoding of genomic databases to succeed there need to be clear benefits to those databases and/or their user community. Similarly, if GBIF is to play a role in this process, we should ask that benefits there would be to GBIF and its users. For INSDC users the benefits are an increasing number of geocoded sequences, making the genomic data better suited for applications such as phylogeography and epidemiology. The geocoded sequences could be uploaded to GBIF, which assigns DOIs to datasets, potentially providing an additional source of citations for genomics researchers.

The verifiability of taxonomic identifications in GBIF is a growing concern given the enormous growth of primary biodiversity data that lacks associated voucher specimens (Troudet et al., 2018). Sequence-based records have the advantage that the identity of sequence-based records can potentially be verified or updated using tools such as BLAST. Hence increasing the amount of sequence-based data in GBIF may help contribute to increased taxonomic accuracy of its data.

## Geocoding approaches

We can divide nucleotide sequences in INSDC into several, partially overlapping categories

1. Those that are geocoded already (e.g., European Molecular Biology Laboratory, 2014).
2. Those that have no latitude and longitude but have some locality information recorded in the sequence feature table (e.g., the “country” qualifier).
3. Those that have a voucher specimen that may have been geocoded already.
4. Those for which coordinates have been published in the paper originally publishing the sequence (or associated supplementary information).

Many category 1 sequences come from sources such as the Barcode of Life Data System (BOLD; http://boldsystems.org) (Ratnasingham et al. 2007, 2013) where geographical coordinates are a key part of the data. Category 2 sequences could be geocoded if we have a tool to convert strings describing geographic localities into coordinates. Category 3 sequences can be geocoded if we can match specimen names in INSDC databases with specimen databases. Category 4 sequences could be geocoded by extracting information from either the paper or its supplementary information. Often this information is presented in idiosyncratic forms and is hard to extract (Page, 2010; Tahsin et al. 2014; Weissenbacher et al., 2015). It is likely that a some of this information will be need to be extracted manually, or at least require some degree of manual intervention.

### GBIF as geocoding tool

One approach to geocoding is to have a large database of locality names and associated geographic coordinates (a gazetteer) and simply do a full text search on a user-supplied query. The coordinates of the locality that best match the query string geocodes that string. This is the approach adopted by tools such as Pelias (https://pelias.io). More sophisticated approaches are possible (e.g., Van Erp et al., 2014; Cardoso et al., 2016) but the use of full text search is attractive as it can make use of existing infrastructure. For example, GBIF has over one billion occurrence records, many of which are geocoded. GBIF also supports full text search using Elasticsearch (https://www.elastic.co), so a simple geocoding tool would be to treat GBIF as a community-constructed gazetteer, and search GBIF using the location information associated with a sequence.

As a proof of concept I built such a tool https://lyrical-money.glitch.me (Fig. 1) which takes a string and find a set of most closely matching localities among the specimens in GBIF. Causal investigation suggests this tool has promise, although it needs formal testing. At present it has a simple API, ideally the results of that API would conform to a standard, such the draft GeocodeJSON specification https://github.com/geocoders/geocodejson-spec/tree/master/draft It also represents uncertainty using the point-radius method (Wieczorek et al., 2004), other methods could be explored.

**Fig. 1.**
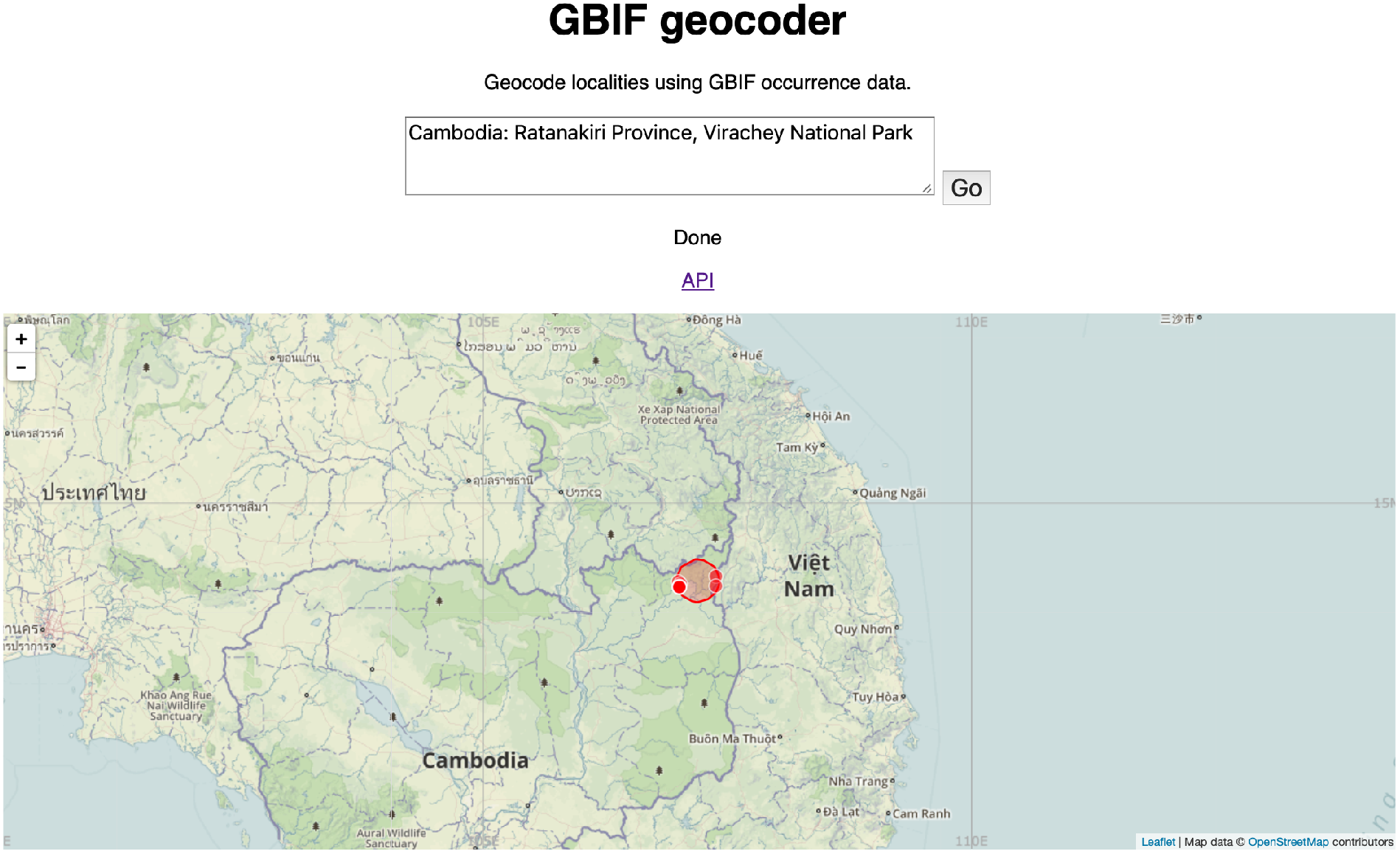
Proof of concept GBIF geocoder, showing the results of geocoding the text “Cambodia: Ratanakiri Province, Virachey National Park”. See https://lyrical-money.glitch.me

### Specimen matching

This approach makes use of the observation that the same specimen may occur in multiple databases (e.g., INSDC, BOLD, GBIF) with various incomplete versions of the associated metadata, such that if those records are combined a more complete record may result (Page, 2016). Specifically, a specimen in GBIF may be geocoded but a sequence from that specimen in INSDC might not be. Linking these two records would geocode the sequence record.

Linking specimens using their codes is problematic as there are multiple ways to refer to the same specimen (Guralnick et al., 2014). I have built a proof of concept tool to match specimen codes to GBIF occurrences https://secretive-rice.glitch.me which endeavours to handle multiple representations of the same specimen code. This tool works best with zoological specimens, which are often cited using a variation of the “Darwin Triple Core” (a combination of the codes representing the institution and collection housing the specimen, together with a catalogue number). Botanical specimens are more often cited using collector codes rather than specimen codes, limiting the utility of the tool I developed. Matching plant specimens will require a different approach.

### Extracting coordinates from the literature and supplementary data

Ideally sequences that are geocoded would have their coordinates included in the sequence database record. However, in numerous cases this information is not included in submissions to the INSDC databases. Instead, it may reside in publications, either in the body of the paper, or in supplementary files. Sometimes those files are in tabular form that can be easily read by machines (e.g., Excel spreadsheets), but often they are in hard to parse formats such as PDFs and Word files. Sometime the data has to be assembled by matching collecting codes in one table to locality codes in another. All of these difficulties hamper automated processing of these sources of information. In many cases the coordinates would have to be extracted manually.

## Data publishing

One issue with any large-scale geocoding of sequences is where that information would best be stored. This has implications for the data publishing process, how the people doing the work are credited, and also how errors are fixed or improvements are incorporated. One approach is to have an intermediate staging point where geocoded sequences are hosted, such that they can be investigated further if necessary. If the output of the geocoding was a Darwin Core Archive file (Wieczorek et al., 2012) then GitHub is an obvious candidate for staging. A Darwin Core Archive comprises one or more comma separated value (CSV) files which can be easily viewed on GitHub, and GitHub repositories can be readily harvested by GBIF. Given the scale of the project the data would likely need to be broken into manageable chunks. One approach would be to use time slices, such as monthly updates to the INSDC databases, perhaps further subdivided by taxonomic group. The associated metadata files files (Fegraus et al. 2005) could cite publication data for all sequences included in each archive, thereby providing provenance for the data and credit for the researchers who did the sequencing. GBIF could harvest these files, adding to its store of geocoded data. INSDC members would need to decide whether they want to update their records with the geocoding, or alternatively treat GBIF as a (LinkOut; https://www.ncbi.nlm.nih.gov/projects/linkout/) partner so that sequences would be linked to the corresponding, geocoded GBIF record.

For Category 4 sequences (geocodes extracted from the literature) we could publish each set of sequences derived from a single paper as a single dataset in GBIF. One approach would be to create a GitHub repository for each paper, adding the text of the paper and any relevant supplementary files and data mining scripts. Once the data have been extracted they could be stored in the repository in CSV files, and the repository packaged in a form suitable to be directly imported into GBIF. The extracted data would get a DOI, which could linked to the DOI for the original publication. By staging the data in Github it would be possible to revisit the data if improved algorithms for text mining became available.

## Conclusion

There is considerable potential for using GBIF to geocode sequence data, either as a gazetter or by cross-linking sequence voucher specimens to GBIF records. These methods could be supplemented by both automated and manual extraction of locality or specimen information from publications. Using GitHub as an intermediate staging platform would enable ready harvesting of the geocoded data by GBIF, as well as easy editing and versioning of data. Implementation of this approach requires the development of several tools and services (such as a GBIF-based geocoded and a specimen matching tool), prototypes of which are already available.

